# Pou4F2/Brn3B Overexpression Promotes the Genesis of Retinal Ganglion Cell-Like Projection Neurons from Late Progenitors

**DOI:** 10.1101/2024.06.08.597922

**Authors:** V.M. Oliveira-Valença, J.M Roberts, V.M. Fernandes-Cerqueira, C.H. Colmerauer, B.C. Toledo, P.L. Santos-França, R. Linden, R.A.P. Martins, M Rocha-Martins, A. Bosco, M.L. Vetter, M.S. Silveira

## Abstract

Retinal ganglion cells (RGCs) are the projection neurons of the retina. In early retinal progenitor cells (RPCs), *Atoh7* orchestrates the developmental RGC program and regulates the expression of critical downstream targets, including *Pou4f* factors. The absence of *Pou4f2* or more POU4F family genes results in defects in RGC differentiation, aberrant axonal elaboration and ultimately RGC death, confirming the requirement of POU4F factors for RGC development and survival, with a critical role in regulating RGC axon outgrowth and pathfinding. Here, we investigated *in vivo* whether ectopic *Pou4f2* expression in late retinal progenitor cells (late RPCs) is sufficient to induce the generation of cells with RGC properties, including projecting axons to the brain. Using a strong ubiquitous promoter to induce *Pou4f2* overexpression in neonates, we observed a change in targeted cell distribution in the retinal tissue, including the presence of cells in the ganglion cell layer and inner plexiform layer with high density of GFP^+^ processes along the retina. Similar results on the induction of neuron processes were obtained when we overexpressed *Pou4f2* in *Atoh7* knockout mice, suggesting that POU4F2 is sufficient to induce them. Single-cell RNA sequencing (scRNA-seq) analysis shows that several RGC-genes (such as *Rbpms*, *Gap*-*43*, *Hs6st3*, and *Foxp2*) are upregulated after *Pou4f2* overexpression. Additionally, gene ontology analysis indicates the induction of genes related to axonogenesis and neuronal differentiation. Imaging throughout the visual pathway revealed high density of axons projecting toward the optic nerve head and extending to brain regions, such as the superior colliculus and geniculate nucleus. Thus, *Pou4f2* induced neurons with specific RGC characteristics that share similarities with resident RGCs and notably project axons that reach brain targets. In conclusion, these results showed that POU4F2 alone was sufficient to promote critical properties of projection neurons from retinal progenitors outside their developmental window.

## INTRODUCTION

Retinal ganglion cells (RGCs) constitute a unique cell population of the retina capable of extending long axons that reach specific brain regions and establish connections. At least 40 subtypes of RGCs have been described based on their transcriptional profile and functions (Rheaume et al., 2018). The ability to project long distance axons is one of the most critical characteristics of RGCs that must be attained in regenerative strategies aimed at restoring vision. Several neuropathies lead to RGC death, such as glaucoma, optic nerve traumas and diabetic retinopathy (Fechtner & Weinreb, 1994; Quigley, 1995; 1999; Halpern & Grosskreutz, 2002; Gupta & Yucel, 2007; Danesh-Meyer & Levin, 2015; Gossman et al., 2016; Killer & Pircher, 2018). RGC degeneration in glaucoma alone is expected to blind 111.8 million people worldwide in 2040 (Tham et al., 2014).

To mitigate the damage from these neuropathies, current research seeks to either decelerate cell loss through RGC neuroprotection or to restore vision via the generation of new RGCs. The potential regenerative cell sources are either manipulated endogenous retinal cells or exogenously generated from embryonic or induced pluripotent stem cells (Gossman et al., 2016; Tanaka et al., 2016; Chao et al., 2017; Oliveira-Valenca et al., 2020; Luo & Chang, 2023; Sharma et al., 2023; Soucy et al., 2023; Tribble et al., 2023). Müller glia (MG), the main endogenous cell source currently under investigation, share epigenetic and transcriptional profiles with the late retinal progenitor cells (late RPCs) (Blackshaw et al., 2004; Ooto et al., 2004; Roesch et al., 2008; Dvoriantchikova et al., 2019).

During retinal development in rodents, late RPCs give rise to most rod photoreceptors (RPs), and also to bipolar cells (BCs), subpopulations of amacrine cells (ACs), and MG in the postnatal period. On the other hand, early RPCs generate RGCs, horizontal cells (HCs), some ACs and cone photoreceptors (CPs) during the embryonic period (Finlay et al., 1989; Saha et al., 1992; Hu et al., 1999; Chow et al., 2001; Bassett & Wallace, 2012; Clark et al., 2019).

Specific molecular programs are activated and restricted to establish the competence of retinal progenitors (RPCs) to differentiate into distinct cell populations and build the mature, functional retina (Young, 1985; Rapaport et al., 2004; Heavner & Pevny, 2012; Centanin & Wittbrodt, 2014; Zechner et al., 2020). Thus, any strategy designed to regenerate functional RGCs will have to reactivate key steps of this gene regulatory network (GRN).

*Atoh7* is an essential competence factor that induces the RGC molecular program (Yang et al., 2003; Mu et al., 2005; Le et al., 2006; Yao et al., 2007; Gao et al., 2014; Maurer et al., 2018; Wu et al., 2021). Crucial downstream effectors of *Atoh7* in the RGC gene regulatory network are the POU domain, class 4, transcriptional factors (POU4F1-3) (Liu et al., 2001; Mu et al., 2005). Several studies have demonstrated that the absence of POU4F2 compromises axonogenesis, as RGCs exhibit aberrant projection along the retina and a thinner or absent optic nerve. Additionally, knockout experiments have shown that even when some RGCs differentiate, they eventually die due to failure to stabilize connections to the brain, thereby demonstrating the role of *Pou4f2* in axonogenesis and survival (Erkman et al., 1996; Gan et al., 1996; Xiang et al., 1996; Gan et al., 1999; Erkman et al., 2000). Some studies also suggest that *Pou4f2* could have a role in RGC specification by suppressing genes required for alternative cell fates including amacrine cells and cones (Qiu et al., 2008; Feng et al., 2011). Shi et al. (2013) demonstrated that besides repressing some cell fates, *Pou4f2* is important to induce the generation of POU4F1^+^/3^+^ and melanopsin RGC subtypes. Moreover, Rasheed et al. (2014) show that *Pou4f2* is expressed in two distinct waves regulated by miRNAs during retinal development and theorize that in early stages it would play a role in RGC specification and, later, in survival and axon guidance.

POU4F2 does not solely work with POU4F members but can also act in conjunction with ISL1 to promote RGC differentiation (Pan et al., 2008; Li et al., 2014; Wu et al., 2015), as well as with DLX1/2 (de Melo et al., 2005; Zhang et al., 2017) and EOMES for RGC development and formation of the optic nerve (Mao et al., 2008). Liu et al. (2000) tested the overexpression of *Pou4f2* during the window of RGC generation in chick (thus in the presence of *Atoh7*) and observed an increase in RGC genesis, suggestive of the sufficiency of this factor to induce RGCs. Moreover, Wu et al. (2015) showed that the combination of POU4F2 with ISL1 was enough to induce RGC specification even in the absence of *Atoh7*, demonstrating that this factor could also play a role for RGC specification when combined with other factors. Interestingly, Todd et al. (2022) demonstrated the *in vivo* conversion of murine Müller glia into RGC-like cells after the combined overexpression of *Ascl1*, *Pou4f2* and *Isl1*. These neurons acquired an immature RGC transcription profile, similar to RGCs in the beginning of their specification, however they did not observe axons in the optic nerve.

Given that *Pou4f2* plays essential roles in RGC development and axonogenesis, our goal in this study was to test whether its overexpression as a single transcription factor in late RPCs could reactivate aspects of the RGC molecular program relevant for RGC differentiation, especially axon growth and pathfinding, outside of their development window. We show that *Pou4f2* overexpression indeed induces the generation of RGC-like cells from late RPCs, which migrate to the inner retina and project axons toward the optic nerve head, along the entire visual pathway and to multiple brain targets. Single cell RNA sequencing (scRNA-seq) reveals the generation of cells expressing RGC genes upon *Pou4f2* overexpression, including genes involved in axonogenesis.

Overall, these data demonstrate that overexpression of *Pou4f2* is sufficient to induce the expression of several RGC features in late progenitors, remarkably axonogenesis, as projections innervate appropriate central visual targets in adult mice.

## MATERIALS AND METHODS

### Experimental animals

All experiments were performed according to international rules and were approved by the Ethics Committee on Animal Experimentation of the Health Sciences Center from Federal University of Rio de Janeiro (CEUA/CCS/UFRJ) and within the guidelines of the Institutional Animal Care and Use Committee of the University of Utah. Both sexes of neonatal (P0, postnatal day 0) Lister hooded rats (RRID:RGC 2312466), C57Bl6/J mice (RRID:IMSR_JAX:000664) and B6J.129-Atoh7^tm1Gla/Mmucd^ (MGI2159015, mentioned here as Atoh7KO), were used for *in vivo* electroporation and analyzed 10 or 30 days post electroporation.

### Plasmids

Plasmids pUb::CST (pUb- human ubiquitin C promoter) and pUb::GFP (Matsuda & Cepko, 2004) were kindly provided by Dr. Michael A. Dyer (St. Jude Hospital, Tennessee, USA). POU4F2 sequence was obtained from CMV::Brn3b plasmid (CAT#:MR223071, Origene) using the restriction enzymes NotI and EcoRI. This fragment was inserted into pUb::GFP vector (previously digested with NotI and Age1) to generate the pUb::Pou4f2 using the oligonucleotide sequences: 5’- GGCCGCTGTAAGCCGTAACGACTAACTCTGC – 3’ and 5’ - CGACATTCGGCATTGCTGATTGAGACGAGCT -3’ to introduce appropriate cut sites. The final concentration of plasmid solution was 5 µg/µL either for control (pUb::CST 70% pUb::GFP 30%) or POU4F2 (pUb::Pou4f2 70% pUb::GFP 30%) experimental groups.

### In vivo electroporation and EdU incorporation

This procedure was performed as described previously (Matsuda & Cepko, 2007; Rocha-Martins et al., 2019). Briefly, P0 Lister hooded rats or mice were anesthetized by hypothermia and the plasmid solution was injected into the subretinal space with a Hamilton syringe with 33G blunt end needle (1 µL or 0.5 µL, respectively to rats or mice, of a 5.0 µg/µL solution containing 0.1% Fast Green Dye (cat# 14335, Cayman Chemical)). Five 99V pulses were applied for 50 ms with 950 ms intervals using a forceps-type electrode (Nepagene, CUY650P7) with Neurgel (Spes Medica). After temperature recovery, pups were returned to the original cage. To avoid variability, both control and experimental groups were from the same litter. Pups received EdU by subcutaneous injection (10μl 2.5mM EdU/gram of body weight, Invitrogen #C10637) at P0 and P1 after the electroporation procedure.

### Immunofluorescence

Protocols used were described previously (Rocha-Martins et al., 2019). Eyes were fixed by immersion in 4% paraformaldehyde in phosphate buffered saline (PBS) overnight, cryoprotected in 30% sucrose in PB, and then immersed in OCT for the preparation of transversal cryosections (10µm) that were mounted on microscope slides treated with 6% silane (cat#440140, Sigma). Electroporated retinas, optic nerves (14µm) and brain cryosections (35µm) were immunoreacted with different primary antibodies (RBPMS, #1830-RBPMS/Phosphosolution, Lot#NB1117g; CHX10, X1180P/Exalpha, Lot#13877; SOX9, AB5535/Milipore, Lot#3063352; NEUN/RBFOX3/Milipore, MAB377, Lot#2453249; RHO, ab98887/Abcam; GAP-43, sc-1086/Santa Cruz) combined with anti-GFP (mAB3E6/Invitrogen Lot#2094035 or A11122/ Invitrogen Lot# 2273763). For the analyses of wholemount retinas, the eyes were fixed overnight in 4% paraformaldehyde and retinas dissected and immersed in 0.5% Triton X-100 for at least 30 minutes. The retinas were then incubated in blocking solution (5% horse serum in 2% Triton X-100) for 1 hour and incubated with primary antibodies for at least 2 days, with slow shaking at 4°C. EdU labelling was performed following the guidelines of the kit (Click-iT™ EdU Cell Proliferation Kit for Imaging, Alexa Fluor™ 647 dye, C10340, Invitrogen).

### Image acquisition

For transverse sections from rat retinas immunostained with antibodies for GFP and specific cell markers, images were acquired using a structural illumination microscope (Imager M2, ApoTome, Zeiss) with Plan-Apochromat 20x/0.8 objective and AxioCam MRm camera. EdU and GAP43 labelling, as well as mouse brains were imaged using a 20x/0.8 by confocal microscopy (Nikon A1, Eclipse Ti2). Wholemount retinas were imaged using oil 40x/1.15 objective by confocal microscopy (Leica TCS-SPE with an AOBS system). Image acquisition settings were maintained across replicate experiments. Software used for image acquisition were Zen Blue, LasX and NIS-Elements. Image processing included the use of maximum intensity projection (MIP) tool and color change and adjustments in brightness in Image J/Fiji. GAP-43 insets (**Figure 2E**) represent one slice of 0.8 µm thickness and the main image was generated using the MIP tool. In the wholemount analysis from Atoh7KO mice (**Figure 3**), we defined the layers using the DAPI staining (and looking at the distance between two nuclei) followed by MIP processing. Images were cut and rotated in Adobe illustrator, which was also used for the manual reconstruction of the optic nerve and optic chiasm from images acquired by structured illumination microscopy.

### Quantitative and statistical analyses

Representative images from retinal cryosections were selected for quantitative analyses. In electroporated retinas, 200 to 500 GFP^+^ cells were counted in all the three nuclear layers for each animal analyzed using Zen Blue software or ImageJ 1.54f. The results were plotted as mean ± SEM with adjusted p value. The statistical analyses were performed using Two-way ANOVA with multiple comparisons and corrected using Sidak’s test (graphs from **Figure 1C, E** and **Figure S1 D**). For the graph in **Figure 1G**, unpaired t-test (two-tailed p value) was performed. Alpha value used was 5.0%. For all analyses, GraphPad Prism software version 8.0.2 was used.

**Figure 1:**
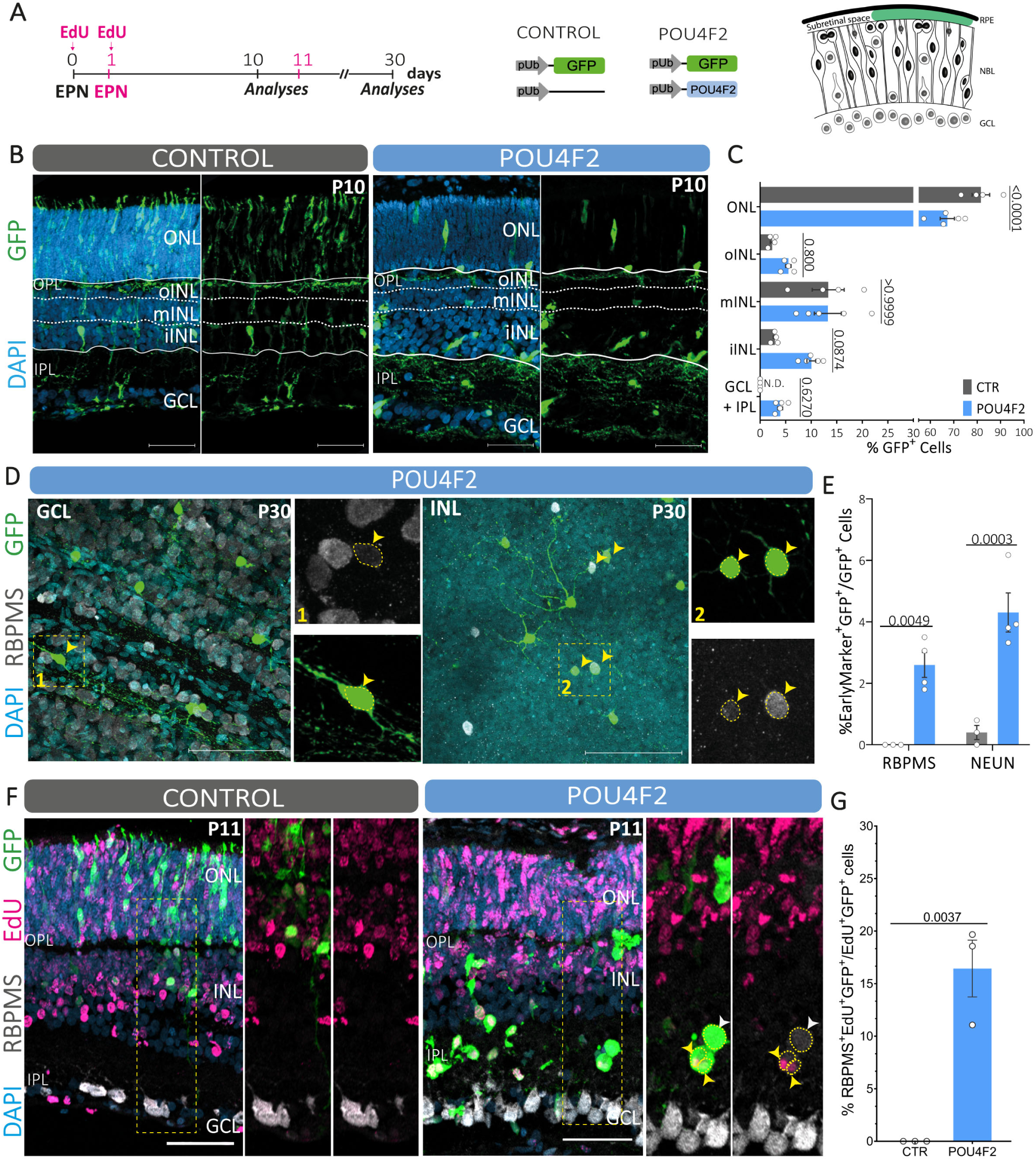
*Pou4f2* promotes the production of RGC-like neurons with complex morphology. **(A)** Experimental design. Rats were electroporated *in vivo* on the date of birth (P0) to induce overexpression of GFP (Control) or Pou4f2 and GFP (Pou4f2), followed by analysis after 10 or 30 days (P10 or P30). For EdU labeling of proliferating cells, EdU injections were performed at P0 and P1, followed by eye electroporation at P1 and harvest after 10 days (design in pink). **(B)** Representative images of radial retinal sections of Control and Pou4f2 conditions after 10 days. **(C)** Quantification of the percentage of GFP^+^ cells at each retinal cell layer (2way ANOVA with Sidak’s test n=4-5). (D) Retinal wholemounts from 30 days after *Pou4f2* electroporation showed GFP^+^RBPMS^+^ cells (yellow arrowhead) located at the GCL (left panel) and INL (right panel). Morphology of some *Pou4f2*-induced neurons is complex with abundant dendrites and an axon. **(E)** Proportion of RBPMS^+^ (RGC marker) and NEUN^+^ (RGC and AC marker) cells in the electroporated population analyzed after 10 days (2way ANOVA with Sidak’s test, n=3-4). **(F)** Radial retinal sections showing EdU labelling used to trace late RPCs and their progeny 10 days post electroporation. Some RBPMS^+^GFP^+^ cells generated upon *Pou4f2* overexpression were EdU^+^ (yellow arrowhead), while others were RBPMS^+^GFP^+^EdU^-^ (white arrowhead). **(G)** Proportion of RBPMS^+^GFP^+^EdU^+^ cells among cycling, electroporated cells (GFP^+^EdU^+^) (unpaired t-test, two-tailed, n=3). For all bar plots each dot represents one animal, bars are mean ±SEM and p-values are shown above the bars. EPN – electroporation; ONL – Outer Nuclear Layer; OPL – Outer Plexiform Layer; oINL-Inner nuclear layer, outer region; mINL-Inner nuclear layer, middle region; iINL - Inner nuclear layer, inner region; IPL – Inner Plexiform Layer; GCL – Ganglion Cell Layer; NFL – Nerve Fiber Layer. Scale bar 50µm.

### scRNA-seq sample preparation

Wild type mice at P0 or P1 were electroporated with the plasmid preparations Pou4f2-GFP or GFP alone injected into the subretinal space. Ten days later, retinas were collected for dissociation. Electroporated retinas were dissociated using papain as described before in Justin Brodie-Kommit et al. (2021). Briefly, papain solution (100 µL papain – 10mg/mL – containing 110 µL of 50mM L-cysteine, 110 µL of 10mM EDTA, 10 µL of 60mM 2-Mercaptoethanol) was pre-incubated at 37°C for 30 minutes before the dissociation. After that, two retinas were added to the papain solution and incubated for 10 minutes at 37°C. 50 µL of DNAse (1mg/mL) was added after 5 minutes of dissociation. To block the reaction, 2.5 mL of neurobasal supplemented with 10% FBS and 500 µL of ovomucoid (10mg/mL) were added. Tubes were spun down for 5min, 300RCF at 4°C and resuspended in 500 µL of neurobasal+10% FBS. Single, viable, GFP^+^ cells were isolated by cell sorting with BD FACSAria 4 laser instrument using 70µm nozzle size and collected into 20 µL DMEM + 5% FBS. Multiple retinas were combined as necessary to obtain 30k GFP^+^ cells for each condition.

### scRNA-seq Library prep and Sequencing

Library generation and sequencing were performed by the Huntsman Cancer Institute High-Throughput Genomics and Bioinformatics Analysis Shared Resource. Sorted cells were prepared with the 10X Genomics Next GEM Single Cell 3’ Gene Expression Library prep v3.1 with UDI. The library was chemically denatured and applied to an Illumina NovaSeq flow cell using the NovaSeq 150x150 bp XP workflow (20043131). Following transfer of the flow cell to an Illumina NovaSeq 6000 instrument, a 150 x 150 cycle paired end sequence run was performed using a NovaSeq 6000 S4 reagent Kit v1.5 (20028312). Sequencing was performed to a depth of at least 200M paired reads/sample.

### scRNA-seq Analysis

Fastq files were aligned to the mouse transcriptome index for mm10 and GFP using cellranger-7.0.1 with expected cells set to 5000. The final number of cells for each condition were: 7,722 control and 9,187 *Pou4f2* overexpression (Pou4f2 OE). The filtered matrix output from cellranger was imported to Seurat v4.3.0.1 and further filtered for cells with less than 4% mitochondrial transcripts, more than 1,200 genes, and fewer than 25,000 transcripts. The control and Pou4f2OE (OE- overexpression) datasets were merged, and SCTransform v0.3.5 pipeline run with all default settings. Clusters were annotated with known markers for retinal cells and combined as appropriate based on those markers (**Table S1** and **Figure 2**). Cluster markers and differentially expressed genes between the experimental conditions were defined using the FindAllMarkers and FindMarkers functions respectively, using default settings (**Tables S4** and **S5**). Integration with developmental data was achieved using Clark et al. Specifically, the developmental dataset was subset for only those cells used for pseudotime analysis, as defined in the cell metadata, and merged with the Pou4f2 single cell dataset. These were combined with the Seurat v5.1.0 IntegrateLayers function using CCA integration (**Figure 4**).

**Figure 2:**
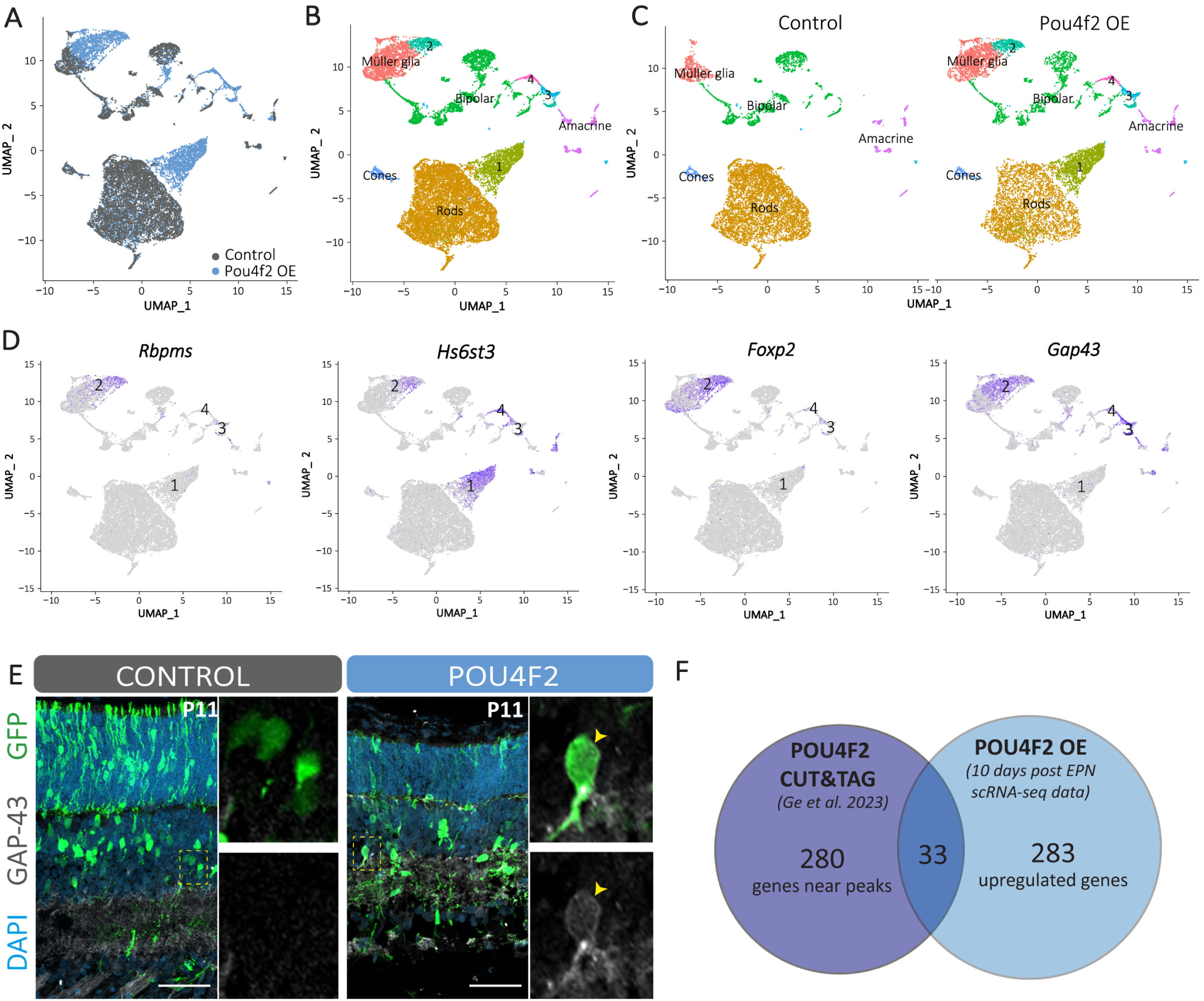
scRNA-seq analysis reveals that *Pou4f2* overexpression induces several RGC-genes. Neonate mice were electroporated *in vivo* (P0) with Gfp (Control) or Pou4f2+Gfp (Pou4f2OE), and after 10 days, retinas were dissociated, GFP^+^ cells collected by FACS, and analyzed by scRNA-seq. **(A)** UMAP plot colored to distinguish control cells (gray) from Pou4f2OE cells (light blue). **(B)** UMAP graph colored by cluster. **(C)** Split UMAPs showing that the new clusters 1-4 are specific for Pou4f2 OE retinas. **(D)** UMAP graphs showing the expression of four RGC-genes (*Rbpms, Hs6st3, Foxp2 and GAP43*). Purple indicates high gene expression levels and gray the absence of gene expression. **(E)** Immunostaining for GAP-43 validates the scRNA-seq result of *Pou4f2* inducing the expression of this gene in some GFP^+^ cells (yellow arrowhead), which is not observed in control condition. **(F)** Intersection between upregulated genes from Pou4f2 OE scRNA-seq (light blue) and previous published CUT&TAG for POU4F2 targets (purple) (Ge et al., 2023).

**Figure 3:**
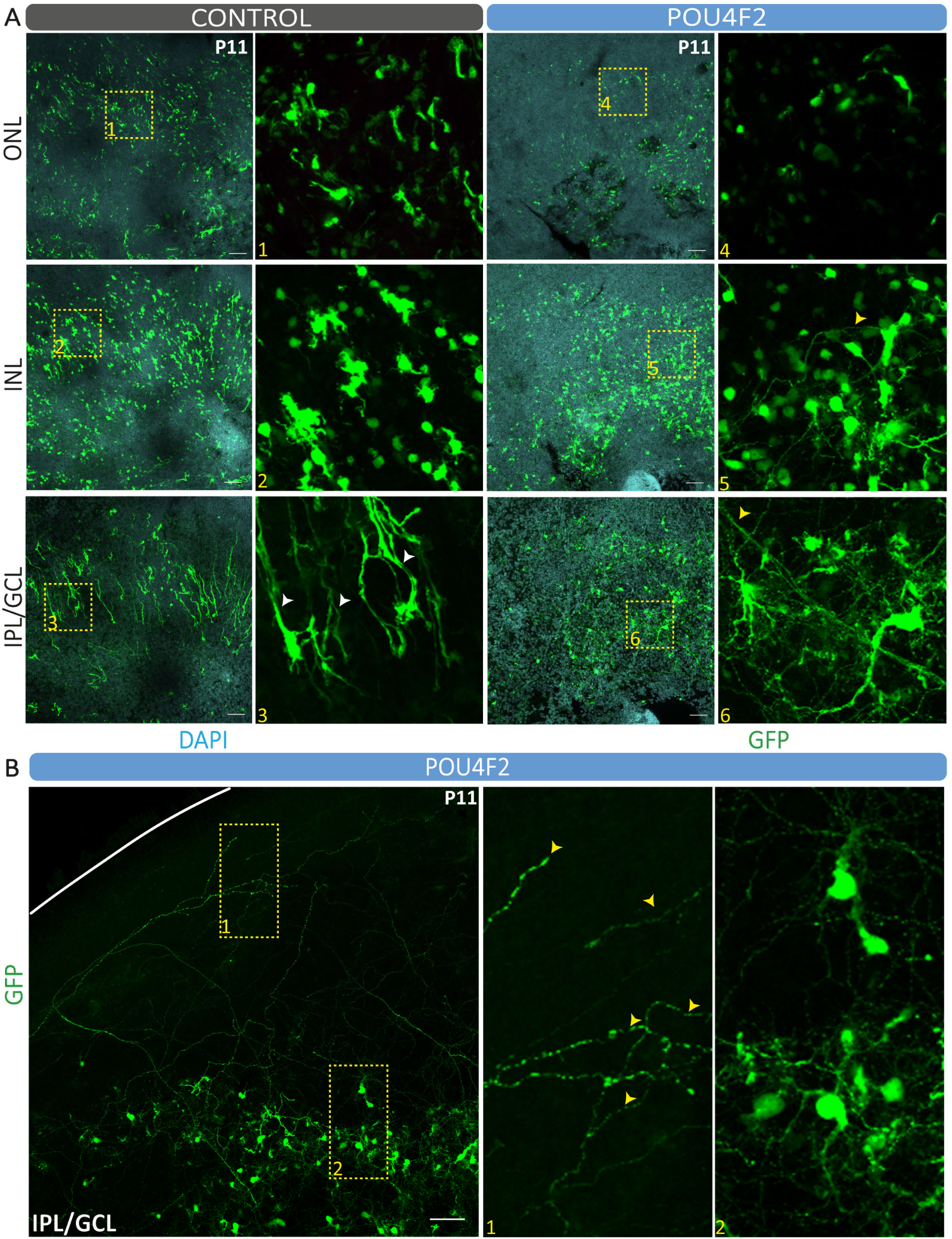
Aberrant projections are detected after *Pou4f2* overexpression in the Atoh7KO mice. Neonate Atoh7KO mice were electroporated *in vivo* (P1) with Gfp (Control; n=2) or Pou4f2+Gfp (POU4F2OE; n=2), and after 10 days, retinas were immunostained for GFP (green). **(A)** Wholemount retinas from each condition (control in the left and Pou4f2OE in the right panel) show electroporated cells (GFP^+^) distributed in the retinal layers: ONL (first line), INL (middle line) and IPL+GCL (third line). GFP+ cells are found in the IPL+GLC (A inset 6; yellow arrowhead indicates the putative axon) while only MG endfeet are detected in Control **(A inset 3, white arrowheads). (B)** After *Pou4f2* overexpression random GFP^+^ axons (yellow arrowheads) are found in the periphery of the retina. The same is not observed in the control condition. ONL – Outer Nuclear Layer; INL-Inner nuclear layer; IPL – Inner Plexiform Layer; GCL – Ganglion Cell Layer. Scale bar 50µm.

**Figure 4:**
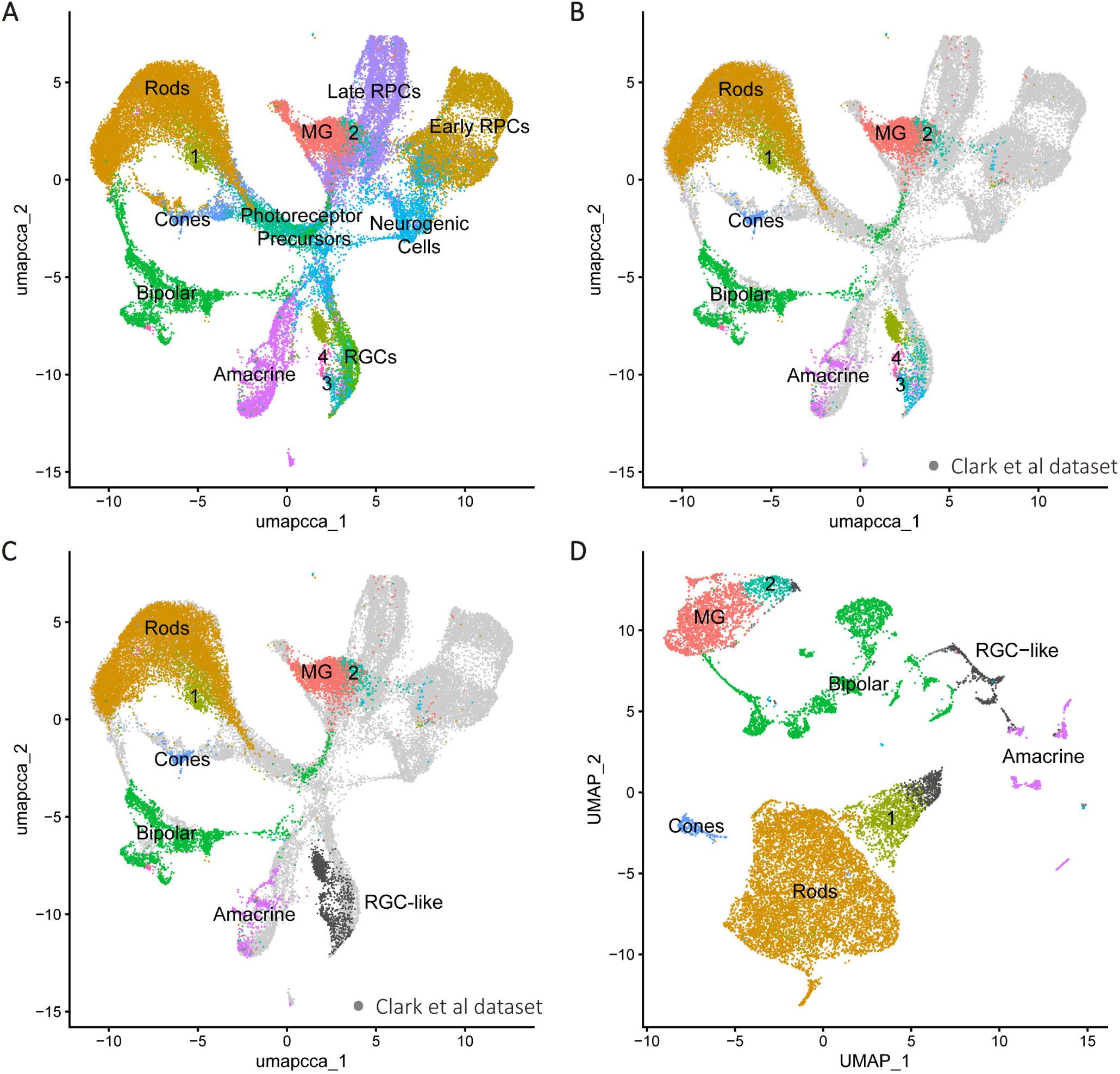
RGC-like cells are clustered near endogenous RGCs. ScRNA-seq dataset from CTR and Pou4f2 overexpression conditions were integrated with published data from E11-P14 retinas (Clark et al., 2019). **(A)** UMAP graph showing both datasets integrated. Clusters 1-4 are the new clusters identified after *Pou4f2* overexpression. **(B)** UMAP graph showing cells from the dataset generated in this paper (colored cells), overlapped with Clark et al (2019) dataset (gray). **(C)** Recoloring cells that are closest to the RGC cluster in dark gray reveals that cells from clusters 1-4 are found in this group. **(D)** Plotting of these RGC-like cells in the UMAP generated from our dataset, using the same colors as in **C**, clearly identifies subpopulations in clusters 1 and 2.

### Gene Set Enrichment Analysis

Gene set enrichment analysis was run on the scRNA-seq Pou4f2 OE dataset using gene sets obtained from the Molecular Signatures Database. Specifically, we used the mouse-M5 ontology gene sets v2023.2. All detectable genes were ranked by average log2 fold change between all control and Pou4f2 OE cells and analyzed with fgsea v1.27.0 using default settings. The results from this analysis are in **Table S3** and **Figure 5**.

**Figure 5:**
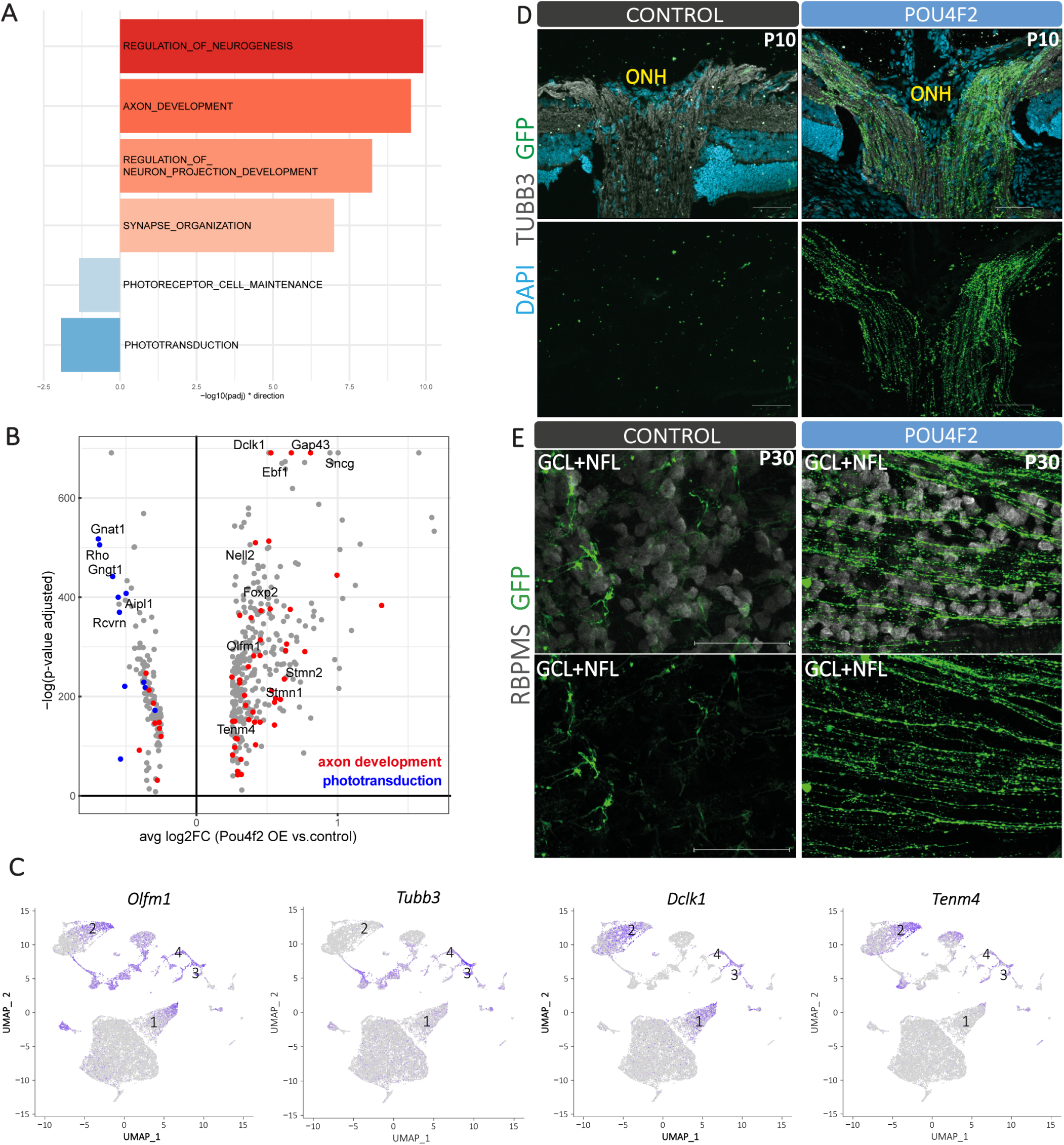
*Pou4f2* overexpression induces axonogenesis. Retinas from mice **(A-C)** or rats **(D-E)** were electroporated *in vivo* on the date of birth (P0) for the overexpression of Gfp (Control) or Pou4f2+Gfp (Pou4f2 OE). After 10 and 30 days, tissues were harvested for dissociation for scRNA-seq or immunostaining. **(A)** Gene ontology analysis of scRNA-seq data reveals the *Pou4f2* induced (in red to orange) and downregulated (in blue) processes. **(B)** Volcano plot shows the distribution of differentially regulated genes in scRNA-seq. Genes of phototransduction (blue dots) and axon development (red dots) are highlighted. **(C)** UMAPs show the expression of some axonogenesis regulators (*Olfm1*, *Tubb3*, *Dclk1* and *Tenm4*) in the *Pou4f2*-induced clusters. Purple indicates high gene expression levels and gray the absence of gene expression. Representative images from optic nerve head **(D)** and wholemounted retinas **(E)** show a high density of GFP^+^ projections in the Pou4f2 OE retina. Scale bar 50 µm **(D)** and 100 µm **(E)**. ONH- optic nerve head. GCL – Ganglion Cell Layer, NFL – Nerve Fiber Layer.

## RESULTS AND DISCUSSION

### Overexpression of *Pou4f2* impacts the fate of late RPCs

The loss of POU4F2 impacts RGC terminal differentiation and survival, through its significant role in axon growth and pathfinding (Erkman et al., 1996; Gan et al., 1996; Xiang et al., 1996; Gan et al., 1999; Erkman et al., 2000). To investigate whether overexpression of *Pou4f2* in late RPCs can promote aspects of RGC differentiation, we performed subretinal *in vivo* electroporation at postnatal day 0 (P0) (Matsuda & Cepko, 2004). (**Figure 1A**). The mature retina is a stratified tissue composed of three nuclear layers where specific cell types are precisely arranged (Dowling, 1987; Livesey & Cepko, 2001; Amini et al., 2017). In line with the known competence of late RPCs, analysis of control retinas 10 days after electroporation showed that the majority of GFP^+^ cells were in the outer nuclear layer (ONL) (81.58 ± 3.76%), fewer were in the inner nuclear layer (INL) (18.56 ± 3.71%), and none to the ganglion cell layer (GCL) (**Figure 1B and 1C**). In comparison, retinas co-electroporated with plasmids for *Pou4f2* and GFP showed fewer GFP^+^ cells at the ONL (67.25 ± 3.04%), a subtle increase in the INL (28.82 ± 3.14%, p=0.0484) and some GFP^+^ cells localized to the ganglion cell layer/inner plexiform layer (GCL-IPL) (3.93 ± 0.47%) (**Figure 1B and C**). These data suggest two major possibilities: i) *Pou4f2* promotes a change in the cell fate of late RPCs, or ii) *Pou4f2* affects the migration of cells derived from late RPCs.

To test the first hypothesis, we performed immunostaining to characterize the identity of GFP^+^ cells. Examining retinas 30 days after *Pou4f2* co-electroporation, we identified GFP^+^ cells which were RBPMS^+^ (a specific RGC marker) in wholemount retinas, while this was not observed after GFP electroporation alone. These RBPMS^+^GFP^+^ cells were in the GCL (left panel) and in the INL (right panel) with some exhibiting complex branching (**Figure 1D**). Quantification 10 days after *Pou4f2* electroporation revealed a significant increase in the proportions of RBPMS^+^ and NEUN^+^ (expressed by both amacrine cells and RGCs) among the GFP^+^ cells relative to controls (**Figure 1E**).

Besides RGC markers, we also immunostained against specific markers for late-born cell types such as rod photoreceptors (RHO^+^), bipolar cells (CHX10^+^) and Müller glia (SOX9^+^), to evaluate whether *Pou4f2* overexpression changes the fate of a specific cell population (**Figure S 1 A-D**). *Pou4f2* significantly reduced the number of RHO^+^GFP^+^ cells whereas no significant changes were found in the numbers of bipolar cells (CHX10^+^) or Müller glia (SOX9^+^) (**Figure S 1 D**). These data suggest that the change in gene expression occurs mostly in progenitors that would normally generate rod photoreceptors.

To confirm whether these cells were derived from late retinal progenitors we performed lineage-tracing experiments through EdU incorporation in cycling cells. EdU was injected at P0 and P1 (followed by electroporation in P1) to label some late RPCs and their daughter cells (**Figure 1A**). Previously, we established that this experimental design resulted in EdU labeling of more than half of GFP^+^ late RPCs and their progeny (62.55 ± 1.97%) after 10 days. Around 16.44 ± 2.70% of this population (GFP^+^EdU^+^) were RBPMS^+^ cells (**Figure 1F and G**) showing that they are newborn cells generated upon *Pou4f2* overexpression. Together, these changes in both gene expression and laminar localization support that *Pou4f2* overexpression induces a cell fate change in late RPC-derived cells.

### Transcriptome analysis reveals induction of RGC and axonogenesis genes

To further characterize the gene expression profile of the *Pou4f2*-induced cells, we isolated GFP-labeled cells by FACS after electroporation with plasmids for GFP alone (CTR) or together with *Pou4f2*, then performed single cell RNA-sequencing (scRNA-seq). Uniform manifold approximation and projection (UMAP) analysis enabled the identification of cells that share transcriptional profiles and can be classified as specific cell types (Becht et al., 2019). Using previously reported cell type specific markers (**Table S1**) (Macosko et al., 2015), we were able to define clusters of late-born cell types among GFP^+^ cells. Similar to the results of our immunostaining experiments (**Figure S1 D**), the majority of control cells were identified as rod photoreceptors (67.2%), a smaller portion as bipolar cells (19.4%), and the minority as Müller glia (6.6%), amacrine cells (4.4%) or cones (1.7%) (**Figure 2B and Figure S1 D and E**). In addition, clusters unique to the *Pou4f2* OE sample were identified (**Figure 2A**, blue dots), indicated as clusters 1, 2, 3 and 4 (**Figure 2B** and **C**). In these *Pou4f2*-clusters (1-4), we detected enrichment of RGC-specific genes, such as *Rbpms* (HÖRNBERG et al., 2013; Rodriguez et al., 2014), *Hs6st3 (Conway et al., 2011)*, *Foxp2* (Rousso et al., 2016; Sato et al., 2017) and *Gap43* (Reh et al., 1987; Kruger et al., 1998) (**Figure 2D**). Moreover, immunostaining for GAP-43 validated scRNA-seq data, showing GAP-43^+^GFP^+^ cells after *Pou4f2* overexpression, but not in control condition (Figure 2E). Again, consistent with our immunostaining the proportions of rod photoreceptors (34.2%) are reduced in favor of novel POU4F2 cell clusters 1-4 (29.4%). There was a reduction in the proportion of bipolar cells (15.5%) and an increase in Müller glia (16.6%) with the caveat that Müller glia and cluster 2 share similarities at the transcriptional level (**Figure S1 E**). This data reinforces that *Pou4f2* overexpression is sufficient to induce changes in the fate of late RPCs, allowing the generation of RGC-like cells from a subset of these progenitors.

Since the role of POU4F2 on RGC specification is incompletely understood (Gan et al., 1996; Xiang, 1998; Rasheed et al., 2014), we hypothesize that several mechanisms may be linked to the genesis of RGC-like cells upon *Pou4f2* overexpression (**Figure 1D** and **Figure 2D**). *Pou4f2* may reprogram a subset of late RPCs through an *Atoh7*-independent pathway (Justin Brodie-Kommit et al., 2021) or induce RGC-related genes in a subset of ATOH7^+^ late RPCs. Indeed, Miesfeld et al. (2018) identified ATOH7^+^ cells in the proliferative layer of the mouse retina at P0. Alternatively, *Pou4f2* may induce RGC characteristics in cells generated from late RPCs through the direct induction of a partial repertoire of RGC genes, such as those responsible for axonogenesis.

To explore these hypotheses, we overexpressed *Pou4f2* in *Atoh7* knockout mice (Atoh7KO) and tested whether the generation of RGC-like cells and projections occur. We first analyzed the cell distribution of GFP^+^ among retinal layers and we found cells located at the ganglion cell layer when *Pou4f2* was overexpressed (**Figure 3A inset 6, B and D**), which is not detected in the control condition (we only detect GFP^+^ MG-end feet, (**Figure 3A inset 3**). When we performed the immunostaining for RBPMS after *Pou4f2* overexpression, even though we could not clearly detect neither GFP^+^RBPMS^+^ nor GFP^-^RBPMS^+^ cells, we identified neurons with complex morphology and aberrant projections which presented a random trajectory or even were directed to the periphery of the retina (**Figure 3A insets 5, 6 and B**). GFP^+^ projections were disorganized probably as a consequence of the absence of axons from the resident RGCs and, consequentially, the optic nerve. These data suggest that, even in the absence of *Atoh7*, *Pou4f2* can induce RGC properties, such as to project axons.

To identify POU4F2-direct targets that could promote changes in the transcriptional profile, we compared differentially expressed genes (DEGs) from our dataset with previously published POU4F2 CUT&TAG data (Ge et al., 2023). Ge et al. identified 313 genes with one or more nearby POU4F2 binding sites. When we compared these putative POU4F2 target genes to the 316 upregulated genes from our scRNA-seq dataset, we observed an intersection of 33 genes (**Figure 2F**), indicating that POU4F2 could have directly induced the expression of these genes. UMAP visualization of the overlapping 33 genes revealed that most of these direct POU4F2-targets were expressed specifically in the new clusters exclusively generated after *Pou4f2* overexpression (**Figure S2** and **Table S2**). Interestingly, many of the overlapping genes are regulators of axon growth, which is consistent with previous studies (Gan et al., 1999; Erkman et al., 2000). These results show that POU4F2 could directly induce changes in the transcriptional profile favoring axonogenesis.

Additionally, to test whether these RGC-like cells were transcriptionally similar to RGCs generated during development, we integrated our scRNA-seq data with a previously published dataset (Clark et al., 2019) encompassing all cell types during retinogenesis (E11-P14). This allowed us to compare the new clusters generated upon *Pou4f2* overexpression with early-born cell types, as well as early and late RPCs (**Figure 4A**). Interestingly, we saw those cells from clusters 3 and 4, as well as some cells from clusters 1 and 2, were close to the RGC cluster in UMAP representation, suggesting that these newly generated cells shared transcriptional similarities with RGCs at a certain level (**Figure 4A and B**).

To represent the distribution of cells from clusters 1-4 that share similarities with RGCs in Clark et al., we recolored these cells as dark gray (**Figure 4C**) and plotted them back onto the first UMAP generated with only our control (CTR) and POU4F2 datasets (**Figure 4D**). This analysis revealed that RGC-like cells from cluster 1 and cluster 2 are positioned at the tip of these clusters, with some cells from cluster 2 closer to cluster 3. The results suggest that *Pou4f2* overexpression was able to induce an RGC-related fate in some cells from clusters 1-4, that are transcriptionally similar to RGCs generated during development.

To define the biological processes induced by *Pou4f2* overexpression, we performed gene ontology (GO) analysis on our scRNA-seq results. This analysis revealed upregulation of genes involved in several neurogenic processes, including axon development, neurogenesis, and synapse organization. Conversely, phototransduction and photoreceptor maintenance genes were downregulated, which could be attributed to the reduction in the population of rod photoreceptors (**Figure 5AB, Table S3**). Further, volcano plot analysis highlights specific genes from the above-regulated biological processes, demonstrating upregulated genes related to axonogenesis (red dots) and downregulated phototransduction genes (blue dots). Notably, RGC-specific markers (SNCG and FOXP2) were significantly upregulated in Pou4f2 OE (**Figure 5B and Table S4**).

UMAP visualization of the gene expression pattern of selected genes linked to axon development by GO analysis, such as *Olfm1* (Nakaya et al., 2008; Nakaya et al., 2012), *Tubb3*, *Dclk1* (Nawabi et al., 2015) and *Tenm4* (Young et al., 2023), showed their elevated expression in the new clusters (1-4) generated by *Pou4f2* overexpression (**Figure 5C**), including *GAP43* (**Figure 2D**). Importantly, 12 of the 33 genes we’ve described as direct POU4F2-targets are in the axon development GO category (Ge et al., 2023) (**Figure S2**).

We also found in our scRNA-seq data other genes that may play important roles in axonogenesis (**Figure 2F** and **Table S2**). For example, *Nav2* (Neuron navigator 2) is responsible for regulating normal cranial nerve development and neurite outgrowth (McNeill et al., 2010) and may play an axonal outgrowth role in RGC-like cells. Additionally, *Ebf1*, Early B-Cell Factor 1, is important for correct axonal topography in the optic chiasm (Jin & Xiang, 2011), in the same way as *Tenm4* (*Ten-m/Odz/teneurin 4*) was described to play a role in binocular circuits, while its absence impacts contralateral projections to specific targets (Young et al., 2023). These genes are upregulated in one or more of the new clusters generated upon *Pou4f2* overexpression (**Figure S2** and **Table S5**). In conclusion, these data show that *Pou4f2* induces genes which are critical for axon growth, and some of them are described as direct targets of POU4F2.

### *Pou4f2*-induced RGC-like cells project axons that reach the brain

To verify if the axonogenesis genes induced by *Pou4f2* would be sufficient for the RGC-like cells to extend axons through the retina, we first analyzed the presence of GFP^+^ processes in retinal transversal sections and wholemounts 10 or 30 days after electroporation, respectively. Images of central retinal sections showed GFP^+^ projections extending to the optic nerve head (ONH) 10 days after *Pou4f2* overexpression (**Figure 5D**). After 30 days, confocal images from retinal wholemounts displayed high density of projections at the GCL+NFL, most of them organized and directed toward the optic nerve head (**Figure 5E**). This result shows that *Pou4f2* overexpression was able to induce axonogenesis not only at the transcriptional but also at the cellular level, with GFP^+^ projections fasciculating together with the axons of resident RGCs.

Previously our group overexpressed *Klf4* in late RPCs and showed the generation of induced RGCs through the activation of an early RGC-program which included the induction of *Eya2 and Atoh7.* However, those neurons were not able to extend axons over long distances. Indeed, we found GFP^+^ projections which reached only the initial segment of the optic nerve, failing to reach the optic chiasm or brain regions (Rocha-Martins et al., 2019).

Image analysis of the rat visual pathway in longitudinal sections 30 days after *Pou4f2* overexpression detected GFP^+^ projections along the entire optic nerve and crossing the optic chiasm (both ipsilateral and contralateral) (**Figure 6A**).

**Figure 6:**
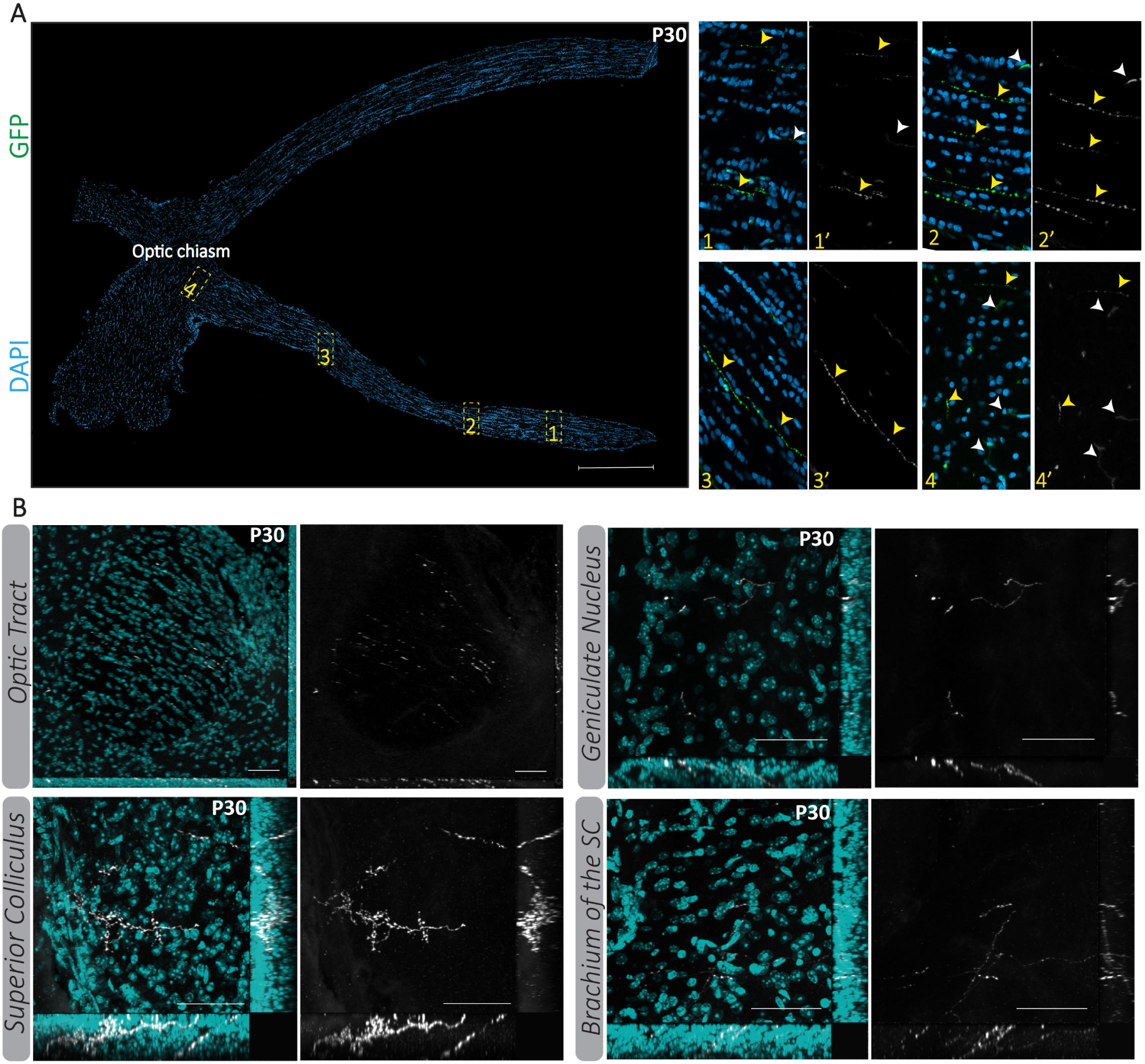
RGC-like cells project eye-to-brain axons. Analysis of GFP^+^ axon extraretinal extension, 30 days after electroporation. **(A)** Optic nerve image reconstruction shows GFP^+^ axons (green, yellow arrowhead) present at the nerve and the chiasm in rats. Insets show high-magnification views at the nerve (1-3) and chiasm (4). White arrowheads indicate non-specific labeling. **(B)** Mouse brain orthogonal images from sagittal brain sections stained for Hoechst, showing GFP^+^ axons (white) at the optic tract, and reaching the superior colliculus and geniculate nucleus. Scale bar 500µm **(A)** and 50µm **(B)**.

Further, to test whether these projections reached brain regions, we analyzed mouse brain sagittal sections 30 days after *Pou4f2* overexpression. We detected GFP^+^ projections in the superior colliculus, branchium of the superior colliculus, optic tract, and geniculate nucleus (**Figure 6B**). All of these regions are known RGC targets in mice (Martersteck et al., 2017) and we did not see any off-target projections.

In conclusion, our study demonstrates that *Pou4f2* overexpression in late RPCs induces the generation of RGC-like cells (RBPMS^+^) located at the GCL and INL, which extend axons toward the optic nerve head, exit the eye, navigate throughout the optic pathway, and reach multiple RGC targets. Additionally, our findings confirm the critical role of POU4F2 in axonogenesis, supported by the identification of POU4F2-direct targets, gene ontology and morphological analyses (**Figure 7**).

**Figure 7:**
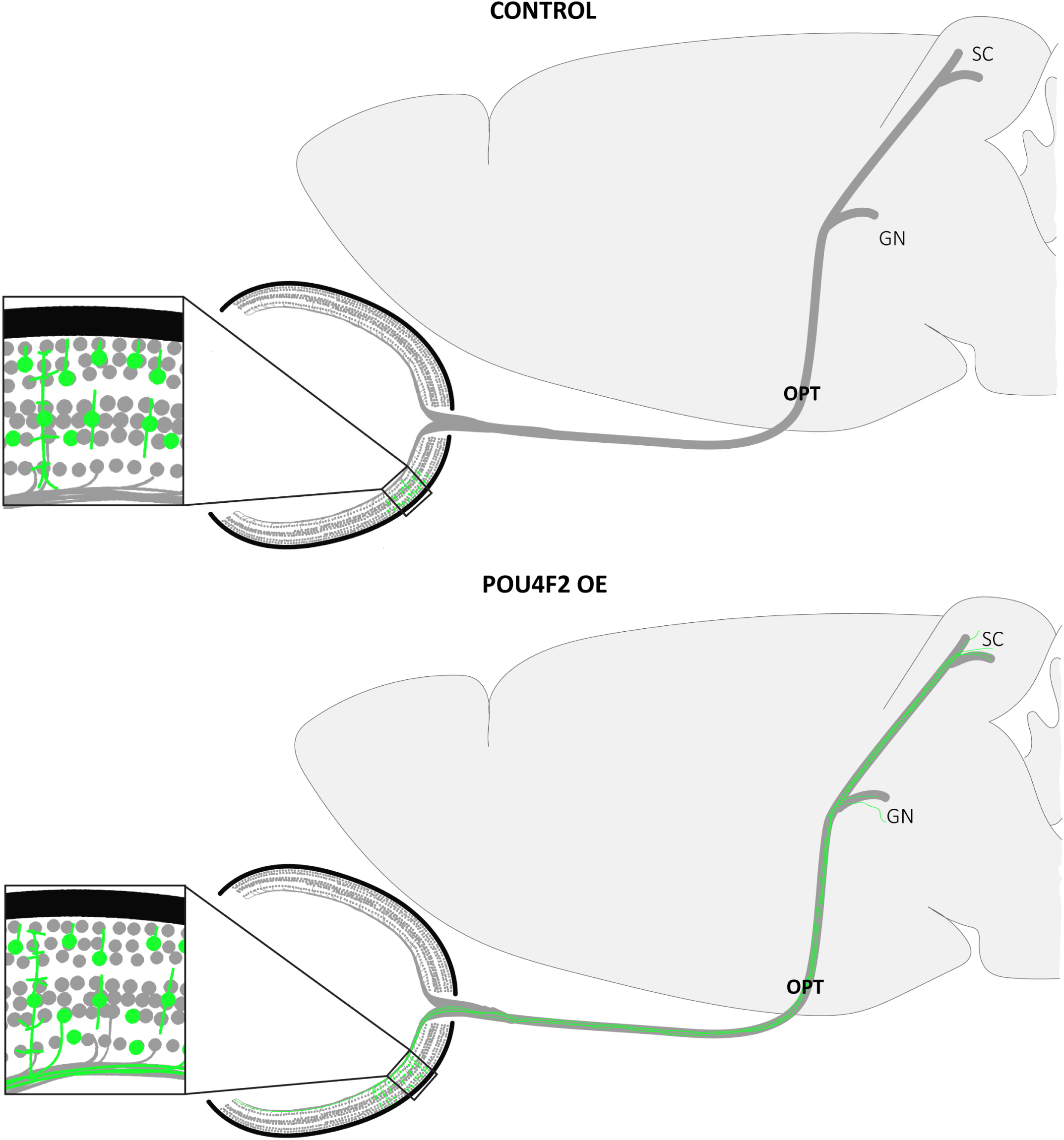
*Pou4f2* induces neurogenesis and axonogenesis of RGC-like cells in a subset of late RPCs. Schematic representation of Control and Pou4f2-overexpressed retinas and visual pathway. In control condition, late RPCs generate rod photoreceptors, bipolar cells, Müller glia and some amacrine cells. *Pou4f2* overexpression induced the generation of RGC-like cells and a high density of projections toward the optic nerve head up to RGC projection areas, such as geniculate nucleus (GN) and superior colliculus (SC). OPT – optic tract.

Since *Pou4f2* overexpression induces genes classified in biological processes related to axonogenesis, we cannot rule out the possibility that in addition to RGC-like cells, other cell types, not expressing RBPMS may have acquired the ability to project axons. This would explain why we identified a high density of projections but a small number of RBPMS^+^GFP^+^ cells, and why in some cases we detect projections at the outer plexiform layer (OPL) (**Figure 1B**) and in the middle of the inner nuclear layer (INL) in transversal sections of the retina. Indeed, some cells from cluster 2 overlap with the RGC cluster (**Figure 4C**) suggesting that they acquired this identity despite maintaining *Sox9* expression, traditionally known as a Müller glia (MG) marker (**Figure S1 E**). Moreover, given that SOX9 is expressed by both MG and late RPCs (Poche et al., 2008), we were unable to ascertain whether the expression of *Sox9* in cluster 2, is attributable to an increase in the number of MG or a persistence of undifferentiated RPCs following *Pou4f2* overexpression. Furthermore, comparative analysis revealed that cells from cluster 2 were near the cluster of late RPC which could indicate a transitional stage without a specific mature cell type identity caused by *Pou4f2* overexpression (**Figure 4B-D**). These data suggest that *Pou4f2* overwrites an axonogenesis program in a subset of neurons allowing them to efficiently extend axons up to the brain.

Our findings partially contrast with a previous study in which *Pou4f2* was overexpressed in late RPCs. In that study, the authors reported the generation of various cell types, including amacrine cells (GABA^+^, GLYT1^+^, PAX6^+^, CALBINDIN^+^), bipolar cells (CHX10^+^), rod photoreceptors (RECOVERIN^+^), Müller glia (GS^+^) and some unidentified cells, in addition to the generation of RGCs (BRN3a^+^). However, they did not highlight any RGC morphological features in the newly generated cells such as the ability to extend projections up to the brain (Qiu et al., 2008). Regarding the differences between the two studies, the authors did not describe the promoter that was used and employed a different strategy for gene delivery (electroporation *vs.* lentivirus). Different levels of induced *Pou4f2* expression in late RPCs may have contributed to the divergent outcomes observed in the two studies.

It is usually expected that in order to regenerate RGCs, certain characteristics need to be achieved, which are: i) specific transcriptional profile; ii) electrophysiological properties; iii) intraretinal connectivity; and iv) ability to project axons to RGC-targets in the brain (Soucy et al., 2023).

In this study, we demonstrated that *Pou4f2* overexpression alone is sufficient to efficiently induce at least one critical property of retinal ganglion cells (RGCs): the ability to extend long-distance axon projections. The RGC-like cells also exhibited transcriptional similarities with RGCs characterized previously (Clark et al., 2019). Importantly, as mentioned before, POU4F2 was able to induce these critical RGC features in an efficient manner, enabling projections to reach brain targets, even if the cells did not possess a complete RGC transcriptional profile. Whether these induced neurons function efficiently as RGCs, receiving and transmitting messages to the brain, remains to be fully elucidated. Nonetheless, the generation of axons suggests a potential avenue for restoring vision through communication between the retinal tissue and the brain.

## Acknowledgements

The authors thank Bernardo Benincá, Franciane de Queiroz Ferreira, Mariana Anjo Barbosa, José Nilson dos Santos, Gildo de Brito Souza, Daianne Torres and José Francisco Tibúrcio for technical support at the Neurogenesis lab, Institute of Biophysics Carlos Chagas Filho, Federal University of Rio de Janeiro (UFRJ) as well as Michelle Chain and Dr. Luiz Dione de Melo from Federal Institute of Rio de Janeiro (IFRJ) for technical support in plasmid validation. We thank Dr. Heitor Affonso de Paula Neto and Dr. Andreza Moreira dos Santos Gama from Unidade Multiusuária de Microscopia Intravital (IMPPG, UFRJ) and Dr. Grasiella Maria Ventura Matioszek from Unidade de Microscopia do Instituto de Ciências Biológicas (UMICB), UFRJ. scRNA-seq data was obtained at the Huntsman Cancer Institute High-Throughput Genomics and Bioinformatics Analysis Shared Resource, University of Utah.

## Authors Contributions

Conceptualization: M.R.-M, M.S.S. and V.M.O.-V; Methodology: V.M.O-V, J.M.R and M.S.S Validation: V.M.O-V and M.S.S.; image analysis: V.M.O.-V, V.M.F-C and C.H.C.; Bioinformatic analysis: J.M.R.; Investigation: V.M.O-V, V.M.F-C, C.H.C, B.C.d.T, P.L.S.-F, M.R-M., J.M.R. and M.S.S.; Writing – review & editing: V.M.O.-V, J.M.R., M.L.V., A.B., P.L.S.-F, R.A.P.M., M.R.M., R.L. and M.S.S.; Supervision: M.S.S.; Project Funding Administration: M.S.S. Funding acquisition: M.S.S., A.B., M.L.V., R.A.P.M. and R.L.

## Declaration of interests

The authors declare no conflict of interests.

## Funding

This work was supported by the International Retinal Research Foundation (IRRF) (MSS and AB). Fellowships for Viviane Oliveira Valença were from Conselho Nacional de Desenvolvimento Científico e Tecnológico (CNPq) (VMO-V: 160968/2018-6 and 200913/2022-0; MSS, MLV and AB: 402078/2022-5); Fundação Carlos Chagas Filho de Amparo à Pesquisa do Estado do Rio de Janeiro (FAPERJ) (VMO-V: E-26/200.214/2020); and Coordenação de Aperfeiçoamento de Pessoal de Nível Superior (CAPES) (V.M.O-V: 88887.512251/2020-00).

## Figure legends

**Figure S1:**
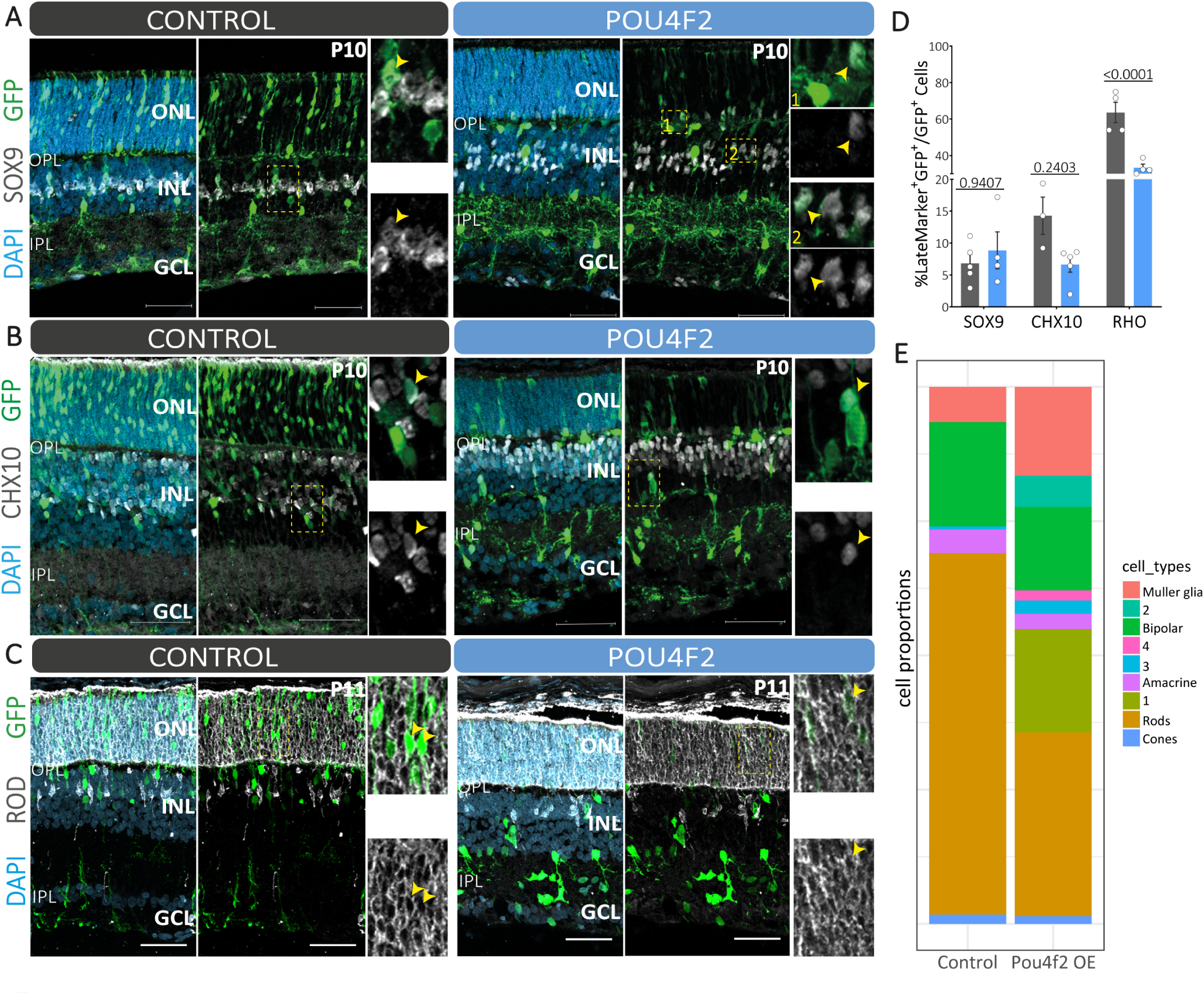
*Pou4f2* reduces the number of rod photoreceptors. Rats were electroporated *in vivo* on the date of birth (P0) to overexpress Gfp (Control) or Pou4f2+Gfp (Pou4f2) and analyzed 10 days later. Images of representative radial retina sections immunostained for: **(A)** SOX9 (Müller glia marker, gray); **(B)** CHX10 (Bipolar cells, gray) and **(C)** Rhodopsin (RHO, rod photoreceptors, grey). Cells with Late Marker^+^GFP^+^ are indicated with a yellow arrowhead in all conditions. **(D)** Quantification of the proportion of LateMarker^+^GFP^+^ cells in the electroporated population (2way ANOVA with Sidak’s test n=3-5). Fewer RHO^+^GFP^+^ cells were detected in POU4F2 condition. **(E)** Graph shows the proportion of each cell type identified in the scRNA-seq analysis between conditions. EPN – electroporation; ONL – Outer Nuclear Layer; OPL – Outer Plexiform Layer; INL-Inner nuclear layer; IPL – Inner Plexiform Layer; GCL – Ganglion Cell Layer. Scale bar 50µm.

**Figure S2:**
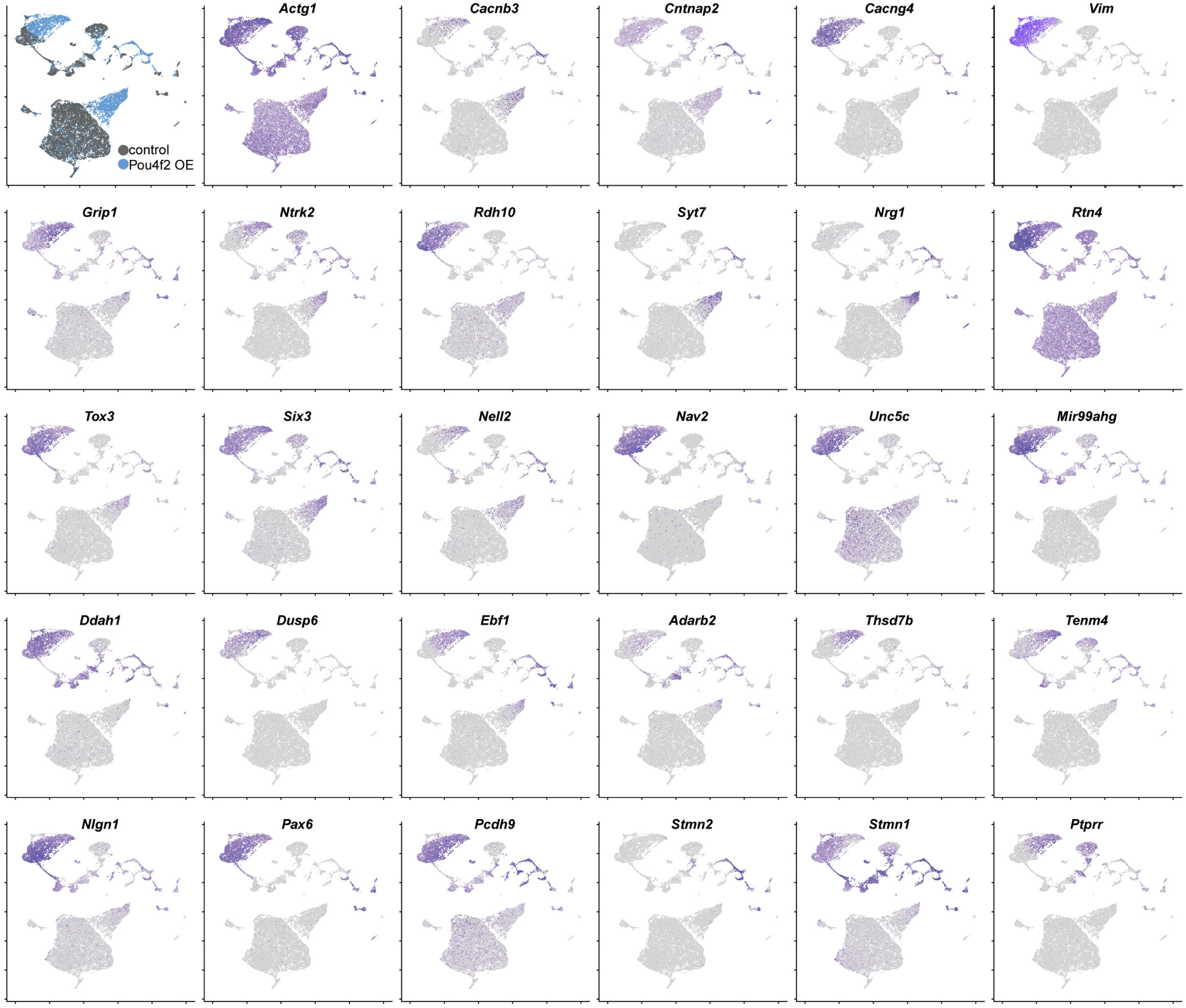
Most POU4F2 direct targets are expressed by the new cell clusters generated after *Pou4f2* overexpression. UMAPs of the 33 genes found in the intersection from the upregulated genes from our scRNA-seq and POU4F2 CUT&TAG data from Ge et al. (2023). Purple indicates high-gene expression levels and gray the absence of gene expression.

## Notes

### Competing Interest Statement

The authors have declared no competing interest.

## References

Amini, R., Rocha-Martins, M., & Norden, C. (2017). Neuronal Migration and Lamination in the Vertebrate Retina. Front Neurosci 11, 742. doi: 10.3389/fnins.2017.00742.

Bassett, E.A., & Wallace, V.A. (2012). Cell fate determination in the vertebrate retina. Trends in neurosciences 35(9), 565–573.

Becht, E., McInnes, L., Healy, J., Dutertre, C.-A., Kwok, I.W.H., Ng, L.G., et al. (2019). Dimensionality reduction for visualizing single-cell data using UMAP. Nature Biotechnology 37(1), 38–44. doi: 10.1038/nbt.4314.

Blackshaw, S., Harpavat, S., Trimarchi, J., Cai, L., Huang, H., Kuo, W.P., et al. (2004). Genomic analysis of mouse retinal development. PLoS biology 2(9), e247.

Centanin, L., & Wittbrodt, J. (2014). Retinal neurogenesis. Development 141(2), 241–244. doi: 10.1242/dev.083642.

Chao, J.R., Lamba, D.A., Klesert, T.R., Torre, A., Hoshino, A., Taylor, R.J., et al. (2017). Transplantation of Human Embryonic Stem Cell-Derived Retinal Cells into the Subretinal Space of a Non-Human Primate. Transl Vis Sci Technol 6(3), 4. doi: 10.1167/tvst.6.3.4.

Chow, R.L., and, & Lang, R.A. (2001). EARLY EYE DEVELOPMENT IN VERTEBRATES. Annu. Rev. Cell Dev. Biol. 17, 255–296.

Clark, B.S., Stein-O’Brien, G.L., Shiau, F., Cannon, G.H., Davis-Marcisak, E., Sherman, T., et al. (2019). Single-Cell RNA-Seq Analysis of Retinal Development Identifies NFI Factors as Regulating Mitotic Exit and Late-Born Cell Specification. Neuron 102(6), 1111–1126.e1115. doi: 10.1016/j.neuron.2019.04.010.

Conway, C.D., Price, D.J., Pratt, T., & Mason, J.O. (2011). Analysis of axon guidance defects at the optic chiasm in heparan sulphate sulphotransferase compound mutant mice. J Anat 219(6), 734–742. doi: 10.1111/j.1469-7580.2011.01432.x.

Danesh-Meyer, H.V., & Levin, L.A. (2015). Glaucoma as a neurodegenerative disease. J Neuroophthalmol 35 Suppl 1, S22–28. doi: 10.1097/WNO.0000000000000293.

de Melo, J., Du, G., Fonseca, M., Gillespie, L.A., Turk, W.J., Rubenstein, J.L., et al. (2005). Dlx1 and Dlx2 function is necessary for terminal differentiation and survival of late-born retinal ganglion cells in the developing mouse retina. Development 132(2), 311–322. doi: 10.1242/dev.01560.

Dowling, J.E. (1987). The retina: an approchable part of the brain. Cambridge and London: The Belknap Press of Harvard University Press.

Dvoriantchikova, G., Seemungal, R.J., & Ivanov, D. (2019). Development and epigenetic plasticity of murine Muller glia. Biochim Biophys Acta Mol Cell Res 1866(10), 1584–1594. doi: 10.1016/j.bbamcr.2019.06.019.

Erkman, L., McEvilly, R.J., Luo, L., Ryan, A.K., Hooshmand, F., O’connell, S.M., et al. (1996). Role of transcription factors a Brn-3.1 and Brn-3.2 in auditory and visual system development. Nature 381(6583), 603–606.

Erkman, L., Yates, P.A., McLaughlin, T., McEvilly, R.J., Whisenhunt, T., O’Connell, S.M., et al. (2000). A POU domain transcription factor–dependent program regulates axon pathfinding in the vertebrate visual system. Neuron 28(3), 779–792.

Fechtner, R.D., & Weinreb, R.N. (1994). Mechanisms of optic nerve damage in primary open angle glaucoma. Survey of ophthalmology 39(1), 23–42.

Feng, L., Eisenstat, D., Chiba, S., Ishizaki, Y., Gan, L., & Shibasaki, K. (2011). Brn-3b inhibits generation of amacrine cells by binding to and negatively regulating DLX1/2 in developing retina. Neuroscience 195, 9–20.

Finlay, B.L., and, & Sengelaub, D.R. (1989). Development of the Vertebrate Retina. Plenum Press • New York and London.

Gan, L., Wang, S.W., Huang, Z., & Klein, W.H. (1999). POU domain factor Brn-3b is essential for retinal ganglion cell differentiation and survival but not for initial cell fate specification. Developmental biology 210(2), 469–480.

Gan, L., Xiang, M., Zhou, L., Wagner, D.S., Klein, W.H., & Nathans, J. (1996). POU domain factor Brn-3b is required for the development of a large set of retinal ganglion cells. Proceedings of the National Academy of Sciences 93(9), 3920–3925.

Gao, Z., Mao, C.A., Pan, P., Mu, X., & Klein, W.H. (2014). Transcriptome of Atoh7 retinal progenitor cells identifies new Atoh7-dependent regulatory genes for retinal ganglion cell formation. Developmental neurobiology 74(11), 1123–1140.

Ge, Y., Chen, X., Nan, N., Bard, J., Wu, F., Yergeau, D., et al. (2023). Key transcription factors influence the epigenetic landscape to regulate retinal cell differentiation. Nucleic Acids Research 51(5), 2151–2176.

Gossman, C.A., Christie, J., Webster, M.K., Linn, D.M., & Linn, C.L. (2016). Neuroprotective Strategies in Glaucoma. Curr Pharm Des 22(14), 2178–2192. doi: 10.2174/1381612822666160128144747.

Gupta, N., & Yucel, Y.H. (2007). Glaucoma as a neurodegenerative disease. Curr Opin Ophthalmol 18(2), 110–114. doi: 10.1097/ICU.0b013e3280895aea.

Halpern, D.L., & Grosskreutz, C.L. (2002). Glaucomatous optic neuropathy: mechanisms of disease. Ophthalmology clinics of North America 15(1), 61–68. doi: 10.1016/s0896-1549(01)00012-8.

Heavner, W., & Pevny, L. (2012). Eye development and retinogenesis. Cold Spring Harb Perspect Biol 4(12). doi: 10.1101/cshperspect.a008391.

Hörnberg, H., Wollerton-Van Horck, F., Maurus, D., Zwart, M., Svoboda, H., Harris, W.A., et al. (2013). RNA-binding protein Hermes/RBPMS inversely affects synapse density and axon arbor formation in retinal ganglion cells in vivo. Journal of Neuroscience 33(25), 10384–10395.

Hu, M., and, & Stephen S. Easter, J. (1999). Retinal Neurogenesis: The Formation of the Initial Central Patch of Postmitotic Cells. Developmental Biology 207, 309–321.

Jin, K., & Xiang, M. (2011). Ebf1 deficiency causes increase of Müller cells in the retina and abnormal topographic projection at the optic chiasm. Biochemical and Biophysical Research Communications 414(3), 539–544. doi: 10.1016/j.bbrc.2011.09.108.

Justin Brodie-Kommit, Brian S. Clark, Qing Shi, Fion Shiau, Dong Won Kim, Jennifer Langel, et al. (2021). Atoh7-independent specification of retinal ganglion cell identity. SCIENCE ADVANCES 7(eabe4983). doi: 10.1126/sciadv.abe4983.

Killer, H., & Pircher, A. (2018). Normal tension glaucoma: review of current understanding and mechanisms of the pathogenesis. Eye 32(5), 924–930.

Kruger, K., Tam, A.S., Lu, C., & Sretavan, D.W. (1998). Retinal ganglion cell axon progression from the optic chiasm to initiate optic tract development requires cell autonomous function of GAP-43. J Neurosci 18(15), 5692–5705. doi: 10.1523/jneurosci.18-15-05692.1998.

Le, T.T., Wroblewski, E., Patel, S., Riesenberg, A.N., & Brown, N.L. (2006). Math5 is required for both early retinal neuron differentiation and cell cycle progression. Developmental biology 295(2), 764–778.

Li, R., Wu, F., Ruonala, R., Sapkota, D., Hu, Z., & Mu, X. (2014). Isl1 and Pou4f2 form a complex to regulate target genes in developing retinal ganglion cells. PLoS One 9(3), e92105. doi: 10.1371/journal.pone.0092105.

Liu, W., Khare, S.L., Liang, X., Peters, M.A., Liu, X., Cepko, C.L., et al. (2000). All Brn3genes can promote retinal ganglion cell differentiation in the chick. Development 127, 3237–3247.

Liu, W., Mo, Z., & Xiang, M. (2001). The Ath5 proneural genes function upstream of Brn3 POU domain transcription factor genes to promote retinal ganglion cell development. Proceedings of the National Academy of Sciences 98(4), 1649–1654.

Livesey, F., & Cepko, C. (2001). Vertebrate neural cell-fate determination: lessons from the retina. Nature Reviews Neuroscience 2(2), 109.

Luo, Z., & Chang, K.-C. (2023). Cell replacement with stem cell-derived retinal ganglion cells from different protocols. Neural Regeneration Research.

Macosko, E.Z., Basu, A., Satija, R., Nemesh, J., Shekhar, K., Goldman, M., et al. (2015). Highly Parallel Genome-wide Expression Profiling of Individual Cells Using Nanoliter Droplets. Cell 161(5), 1202–1214. doi: 10.1016/j.cell.2015.05.002.

Mao, C.-A., Kiyama, T., Pan, P., Furuta, Y., Hadjantonakis, A.-K., & Klein, W.H. (2008). Eomesodermin, a target gene of Pou4f2, is required for retinal ganglion cell and optic nerve development in the mouse. Development 135(2), 271–280. doi: 10.1242/dev.009688.

Martersteck, E.M., Hirokawa, K.E., Evarts, M., Bernard, A., Duan, X., Li, Y., et al. (2017). Diverse Central Projection Patterns of Retinal Ganglion Cells. Cell Rep 18(8), 2058–2072. doi: 10.1016/j.celrep.2017.01.075.

Matsuda, T., & Cepko, C.L. (2004). Electroporation and RNA interference in the rodent retina in vivo and in vitro. Proc Natl Acad Sci U S A 101(1), 16–22. doi: 10.1073/pnas.2235688100 2235688100 [pii].

Matsuda, T., & Cepko, C.L. (2007). Controlled expression of transgenes introduced by in vivo electroporation. Proc Natl Acad Sci U S A 104(3), 1027–1032. doi: 10.1073/pnas.0610155104.

Maurer, K.A., Kowalchuk, A., Shoja-Taheri, F., & Brown, N.L. (2018). Integral bHLH factor regulation of cell cycle exit and RGC differentiation. Developmental Dynamics 247(8), 965–975. doi: 10.1002/dvdy.24638.

McNeill, E.M., Roos, K.P., Moechars, D., & Clagett-Dame, M. (2010). Nav2 is necessary for cranial nerve development and blood pressure regulation. Neural development 5, 1–14.

Miesfeld, J.B., Glaser, T., & Brown, N.L. (2018). The dynamics of native Atoh7 protein expression during mouse retinal histogenesis, revealed with a new antibody. Gene Expr Patterns 27, 114–121. doi: 10.1016/j.gep.2017.11.006.

Mu, X., Fu, X., Sun, H., Beremand, P.D., Thomas, T.L., & Klein, W.H. (2005). A gene network downstream of transcription factor Math5 regulates retinal progenitor cell competence and ganglion cell fate. Developmental Biology 280(2), 467–481.

Nakaya, N., Lee, H.S., Takada, Y., Tzchori, I., & Tomarev, S.I. (2008). Zebrafish olfactomedin 1 regulates retinal axon elongation in vivo and is a modulator of Wnt signaling pathway. J Neurosci 28(31), 7900–7910. doi: 10.1523/jneurosci.0617-08.2008.

Nakaya, N., Sultana, A., Lee, H.S., & Tomarev, S.I. (2012). Olfactomedin 1 interacts with the Nogo A receptor complex to regulate axon growth. J Biol Chem 287(44), 37171–37184. doi: 10.1074/jbc.M112.389916.

Nawabi, H., Belin, S., Cartoni, R., Williams, Philip R., Wang, C., Latremolière, A., et al. (2015). Doublecortin-Like Kinases Promote Neuronal Survival and Induce Growth Cone Reformation via Distinct Mechanisms. Neuron 88(4), 704–719. doi: 10.1016/j.neuron.2015.10.005.

Oliveira-Valenca, V.M., Bosco, A., Vetter, M.L., & Silveira, M.S. (2020). On the Generation and Regeneration of Retinal Ganglion Cells. Front Cell Dev Biol 8, 581136. doi: 10.3389/fcell.2020.581136.

Ooto, S., Akagi, T., Kageyama, R., Akita, J., Mandai, M., Honda, Y., et al. (2004). Potential for neural regeneration after neurotoxic injury in the adult mammalian retina. Proceedings of the National Academy of Sciences of the United States of America 101(37), 13654–13659.

Pan, L., Deng, M., Xie, X., & Gan, L. (2008). ISL1 and BRN3B co-regulate the differentiation of murine retinal ganglion cells. Development 135(11), 1981–1990. doi: 10.1242/dev.010751.

Poche, R.A., Furuta, Y., Chaboissier, M.C., Schedl, A., & Behringer, R.R. (2008). Sox9 is expressed in mouse multipotent retinal progenitor cells and functions in Muller glial cell development. J Comp Neurol 510(3), 237–250. doi: 10.1002/cne.21746.

Qiu, F., Jiang, H., & Xiang, M. (2008). A comprehensive negative regulatory program controlled by Brn3b to ensure ganglion cell specification from multipotential retinal precursors. J Neurosci 28(13), 3392–3403. doi: 10.1523/JNEUROSCI.0043-08.2008.

Quigley, H.A. (1995). Ganglion cell death in glaucoma: pathology recapitulates ontogeny. Australian and New Zealand journal of ophthalmology 23(2), 85–91.

Quigley, H.A. (1999). Neuronal death in glaucoma. Progress in retinal and eye research 18(1), 39–57.

Rapaport, D.H., Wong, L.L., Wood, E.D., Yasumura, D., & LaVail, M.M. (2004). Timing and topography of cell genesis in the rat retina. J Comp Neurol 474(2), 304–324. doi: 10.1002/cne.20134.

Rasheed, V.A., Sreekanth, S., Dhanesh, S.B., Divya, M.S., Divya, T.S., Akhila, P.K., et al. (2014). Developmental wave of Brn3b expression leading to RGC fate specification is synergistically maintained by miR-23a and miR-374. Dev Neurobiol 74(12), 1155–1171. doi: 10.1002/dneu.22191.

Reh, T.A., Redshaw, J.D., & Bisby, M.A. (1987). Axons of the pyramidal tract do not increase their transport of growth-associated proteins after axotomy. Brain Res 388(1), 1–6.

Rheaume, B.A., Jereen, A., Bolisetty, M., Sajid, M.S., Yang, Y., Renna, K., et al. (2018). Single cell transcriptome profiling of retinal ganglion cells identifies cellular subtypes. Nat Commun 9(1), 2759. doi: 10.1038/s41467-018-05134-3.

Rocha-Martins, M.c., de Toledo, B.C., Santos-França, P.L., Oliveira-Valença, V.M., Vieira-Vieira, C.H., Matos-Rodrigues, G.E., et al. (2019). *De novo* genesis of retinal ganglion cells by targeted expression of *Klf4 in vivo*. Development 146(16), dev176586. doi: 10.1242/dev.176586.

Rodriguez, A.R., Müller, S., Pérez, L., & Brecha, N.C. (2014). The RNA binding protein RBPMS is a selective marker of ganglion cells in the mammalian retina. Journal of Comparative Neurology 522(6), 1411–1443.

Roesch, K., Jadhav, A.P., Trimarchi, J.M., Stadler, M.B., Roska, B., Sun, B.B., et al. (2008). The transcriptome of retinal Muller glial cells. J Comp Neurol 509(2), 225–238. doi: 10.1002/cne.21730.

Rousso, David L., Qiao, M., Kagan, Ruth D., Yamagata, M., Palmiter, Richard D., & Sanes, Joshua R. (2016). Two Pairs of ON and OFF Retinal Ganglion Cells Are Defined by Intersectional Patterns of Transcription Factor Expression. Cell Reports 15(9), 1930–1944. doi: 10.1016/j.celrep.2016.04.069.

Saha, M.S., Servetnick, M., and, & Grainger, R.M.1 (1992). Vertebrate eye development. Current Opinion in Genetics and Development 2, 582–588.

Sato, C., Iwai-Takekoshi, L., Ichikawa, Y., & Kawasaki, H. (2017). Cell type-specific expression of FoxP2 in the ferret and mouse retina. Neuroscience Research 117, 1–13. doi: 10.1016/j.neures.2016.11.008.

Sharma, T.P., Ho, K., Bodi, N., & Hameed, S.S. (2023). Transplantation strategy of human retinal ganglion cells for glaucoma. Investigative Ophthalmology & Visual Science 64(8), 3846–3846.

Shi, M., Kumar, S.R., Motajo, O., Kretschmer, F., Mu, X., & Badea, T.C. (2013). Genetic interactions between Brn3 transcription factors in retinal ganglion cell type specification. PloS one 8(10), e76347.

Soucy, J.R., Aguzzi, E.A., Cho, J., Gilhooley, M.J., Keuthan, C., Luo, Z., et al. (2023). Retinal ganglion cell repopulation for vision restoration in optic neuropathy: a roadmap from the RReSTORe Consortium. Molecular Neurodegeneration 18(1), 64. doi: 10.1186/s13024-023-00655-y.

Tanaka, T., Yokoi, T., Tamalu, F., Watanabe, S.-I., Nishina, S., & Azuma, N. (2016). Generation of retinal ganglion cells with functional axons from mouse embryonic stem cells and induced pluripotent stem cells. Investigative ophthalmology & visual science 57(7), 3348–3359.

Tham, Y.-C., Li, X., Wong, T.Y., Quigley, H.A., Aung, T., & Cheng, C.-Y. (2014). Global prevalence of glaucoma and projections of glaucoma burden through 2040: a systematic review and meta-analysis. Ophthalmology 121(11), 2081–2090.

Todd, L., Jenkins, W., Finkbeiner, C., Hooper, M.J., Donaldson, P.C., Pavlou, M., et al. (2022). Reprogramming Muller glia to regenerate ganglion-like cells in adult mouse retina with developmental transcription factors. Sci Adv 8(47), eabq7219. doi: 10.1126/sciadv.abq7219.

Tribble, J.R., Hui, F., Quintero, H., El Hajji, S., Bell, K., Di Polo, A., et al. (2023). Neuroprotection in glaucoma: Mechanisms beyond intraocular pressure lowering. Molecular Aspects of Medicine 92, 101193.

Wu, F., Bard, J.E., Kann, J., Yergeau, D., Sapkota, D., Ge, Y., et al. (2021). Single cell transcriptomics reveals lineage trajectory of retinal ganglion cells in wild-type and Atoh7-null retinas. Nature communications 12(1), 1–20.

Wu, F., Kaczynski, T.J., Sethuramanujam, S., Li, R., Jain, V., Slaughter, M., et al. (2015). Two transcription factors, Pou4f2 and Isl1, are sufficient to specify the retinal ganglion cell fate. Proc Natl Acad Sci U S A 112(13), E1559-1568. doi: 10.1073/pnas.1421535112.

Xiang, M. (1998). Requirement for Brn-3b in early differentiation of postmitotic retinal ganglion cell precursors. Developmental biology 197(2), 155–169.

Xiang, M., Zhoui, L., Macke, J.P., Yoshioka, T., Hendry, S.H.C., Eddy, R.L., et al. (1996). The Brn-3 Family of POU-Domain Factors: Primary Structure, Binding Specificity, and Expression in subset of Retinal Ganglion Cells and Somatosensory Neurons. Journal of Neuroscience 15, 4762–4785.

Yang, Z., Ding, K., Pan, L., Deng, M., & Gan, L. (2003). Math5 determines the competence state of retinal ganglion cell progenitors. Developmental biology 264(1), 240–254.

Yao, J., Sun, X., Wang, Y., Xu, G., & Qian, J. (2007). Math5 promotes retinal ganglion cell expression patterns in retinal progenitor cells. Molecular Vision 13, 1066.

Young, R.W. (1985). Cell differentiation in the retina of the mouse. The Anatomical Record 212(2), 199–205.

Young, T.R., Black, D., Mansuri, H., Oohashi, T., Zhou, X.H., Sawatari, A., et al. (2023). Ten-m4 plays a unique role in the establishment of binocular visual circuits. Dev Neurobiol 83(3-4), 104–124. doi: 10.1002/dneu.22912.

Zechner, C., Nerli, E., & Norden, C. (2020). Stochasticity and determinism in cell fate decisions. Development 147(14). doi: 10.1242/dev.181495.

Zhang, Q., Zagozewski, J., Cheng, S., Dixit, R., Zhang, S., de Melo, J., et al. (2017). Regulation of Brn3b by DLX1 and DLX2 is required for retinal ganglion cell differentiation in the vertebrate retina. Development 144(9), 1698–1711.

